# Systemic Nanobubbles Enable Ultrasound-Guided STING Immunotherapy in Breast Cancer

**DOI:** 10.64898/2026.08.13.744654

**Authors:** Nazia Hafeez, Sina Khorsandi, Ruoqi Gao, Anbarin Khalid, Shariq Ali, Tara Movaghar, Shea Garland, Caroline de Gracia Lux, Jacques Lux

**Affiliations:** Department of Radiology, The University of Texas Southwestern Medical Center, Dallas, Texas 75390, USA; Department of Biomedical Engineering, The University of Texas Southwestern Medical Center, Dallas, Texas 75390, USA

## Abstract

Activation of the STING pathway can induce potent antitumor immunity, but effective delivery of STING agonists to the tumor while limiting systemic exposure remains challenging. We previously developed MUSIC, an ultrasound-guided platform that uses microbubbles (MBs) to deliver the STING agonist 2′3′-cGAMP and locally activate antitumor immunity. However, the vascular confinement of MBs and the need for intratumoral administration limit the potential for systemic tumor targeting. To overcome these limitations, we developed **SONATA** (**S**ystemic **O**ncotherapy using **N**anobubbles for **A**coustically-guided **T**umor **A**ctivation), which employs nanobubbles (NBs) that are approximately 10-fold smaller than conventional MBs, enabling systemic administration and tumor extravasation. Following NB accumulation within tumors, ultrasound exposure triggers localized cGAMP release, facilitating delivery to targeted CD11b^+^ antigen-presenting cells (APCs) and STING activation with spatial and temporal control. NBs are composed of the same components as MBs, including phospholipid shells and a perfluorobutane core and are functionalized with anti-CD11b antibodies to target CD11b^+^ APCs and spermine-modified dextran to stably load cGAMP through nanocomplex formation. Upon ultrasound activation, SONATA induced phosphorylation of STING, TBK1, and IRF3 and increased IFN-β production in bone marrow-derived macrophages. In an orthotopic breast cancer model, intravenously administered SONATA combined with tumor-localized ultrasound significantly inhibited tumor growth compared with controls. Furthermore, SONATA synergized with immune checkpoint blockade prolonged the median survival of tumor-bearing mice. Collectively, these findings establish SONATA as a systemically administered immunotherapy platform that enables ultrasound-guided, spatially controlled STING activation.

## Introduction

Cancer immunotherapy has transformed the treatment landscape for many malignancies, particularly through immune checkpoint inhibitors (ICIs) [1–3]. However, a substantial proportion of patients fail to respond because their tumors remain immunologically “cold,” characterized by limited innate immune activation and poor T-cell infiltration. Activating innate immunity within the tumor microenvironment represents a promising strategy to convert these tumors into immunologically “hot” tumors and enhance the efficacy of ICIs. Antigen-presenting cells (APCs), including macrophages and dendritic cells, are central regulators of this process, as they orchestrate antitumor immunity through antigen presentation and production of pro-inflammatory cytokines that promote tumor-specific T-cell responses [4–5].

Among innate immune pathways, the cyclic GMP-AMP synthase–stimulator of interferon genes (cGAS-STING) pathway has emerged as an attractive therapeutic target due to its ability to induce robust type I interferon (IFN-I) signaling, promote APC activation, enhance antigen presentation, and stimulate adaptive immune responses [8–11]. The endogenous STING ligand 2′3′-cyclic GMP-AMP (2′3′-cGAMP) is a potent cyclic dinucleotide agonist with substantial therapeutic potential. However, its highly anionic and hydrophilic structure limits passive plasma-membrane transport and cytosolic bioavailability, while extracellular degradation, short systemic persistence, and nonspecific distribution further restrict its therapeutic utility [12,13]. In addition, broad or poorly localized STING activation can generate dose-limiting inflammatory toxicity, whereas intratumoral administration of STING agonists may be impractical for deep, disseminated, or inaccessible lesions. Accordingly, biomaterial platforms that achieve cell-selective intracellular delivery and spatiotemporally controlled STING activation are needed to fully realize the therapeutic potential of cGAMP.

To address these challenges, we previously developed **MUSIC** (**M**icrobubble-assisted **U**ltra**S**ound-guided **I**mmunotherapy of **C**ancer), an ultrasound-responsive microbubble platform that enables localized cytosolic delivery of cGAMP to APCs through ultrasound-mediated sonoporation [14,15]. MUSIC elicited potent antitumor immune responses in primary and metastatic murine breast-cancer models and enhanced therapeutic responses to programmed cell death protein 1 (PD-1) blockade. Subsequent work further demonstrated that modified ultrasound-responsive MUSIC microbubbles could enhance immune checkpoint blockade in melanoma. Nevertheless, the micron-scale dimensions of conventional microbubbles limit their extravascular access after intravenous administration, and their use for tumor-directed therapeutic delivery has generally required local or intratumoral administration.

Nanobubbles (NBs) have emerged as a potential alternative to microbubbles because their submicron dimensions may promote prolonged tumor retention and access to the extravascular tumor compartment in permissive tumor vasculature [16–18,24,26]. Preclinical studies have reported NB extravasation, interstitial transport, and enhanced retention relative to microbubbles [16–18,20,21]. In particular, NB size, shell composition, ligand-mediated binding, tumor vascular permeability, and extracellular-matrix architecture influence their capacity for transvascular transport and persistence within tumor tissue [19–22,24]. Ultrasound exposure can additionally induce mechanical bioeffects, including sonoporation and transient increases in vascular and plasma-membrane permeability, thereby enhancing local payload release and cellular uptake. However, the extent of intact-NB extravasation is formulation- and tumor-dependent, and rigorous control of size distribution and microbubble contamination remains essential when interpreting NB pharmacokinetics and acoustic activity.

Here, we engineered **SONATA** (**S**ystemic **O**ncotherapy using **N**anobubbles for **A**coustically-guided **T**umor **A**ctivation), an ultrasound-responsive NB platform designed for systemic delivery of cGAMP and localized activation of the STING pathway. We hypothesized that adapting the APC-targeting and cGAMP-loading strategies established with MUSIC to nanoscale NBs would enable CD11b^+^ APC-targeted, ultrasound-guided cytosolic delivery of cGAMP within tumors after systemic administration while overcoming the predominantly vascular confinement of conventional microbubbles. Following intravenous administration, antibody-conjugated NBs are designed to accumulate within tumors, bind target APCs through CD11b-mediated recognition, and release cGAMP upon localized ultrasound exposure. Ultrasound-mediated NB activation is expected to transiently permeabilize the plasma membrane, facilitating cytosolic cGAMP delivery and reducing reliance on endocytic uptake (Figure 1).

**Figure 1.**
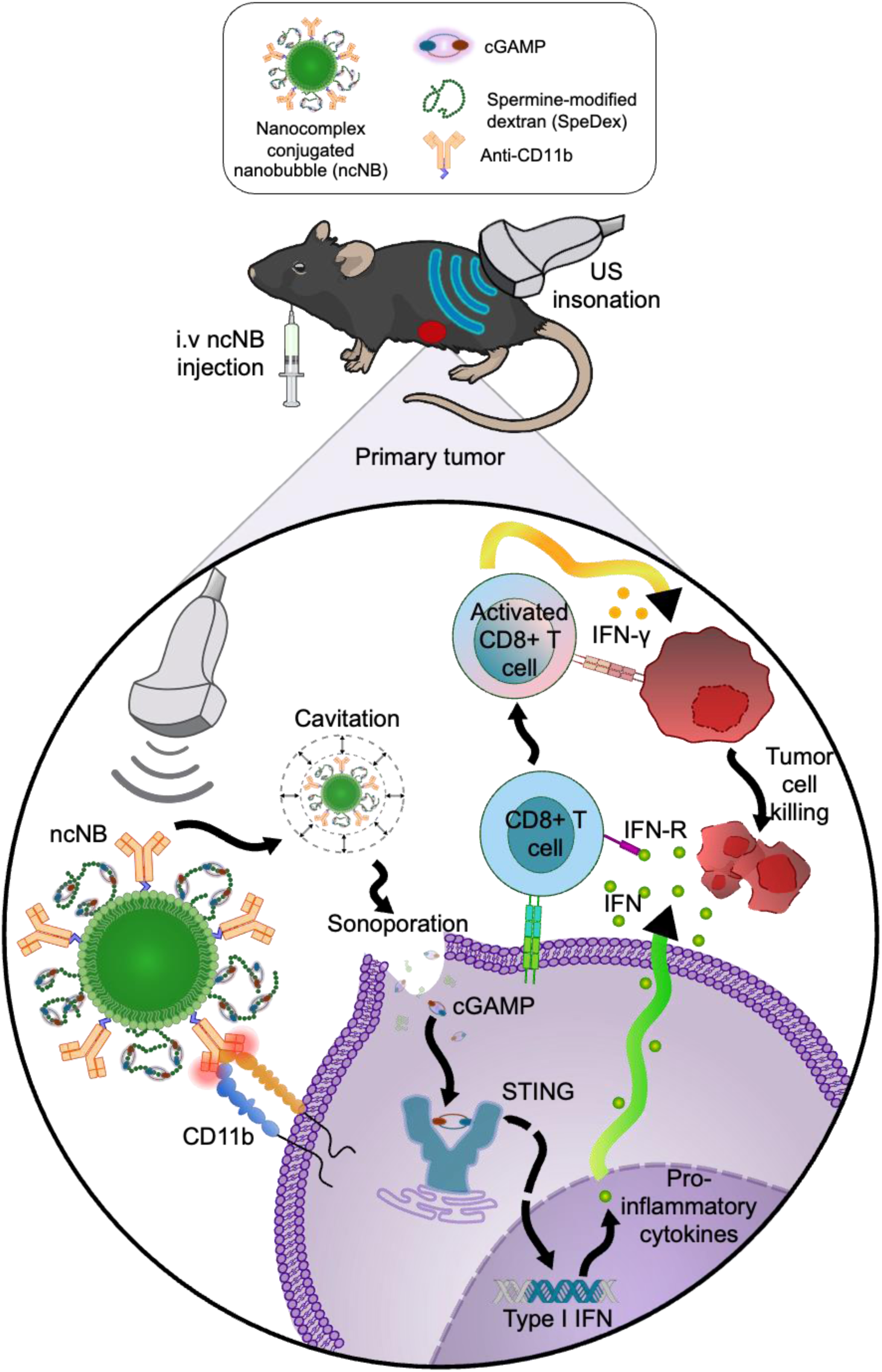
SONATA mechanism of action. Intravenously administered nanocomplex-conjugated nanobubbles (ncNBs) accumulate in the primary tumor and interact with antigen-presenting cells (APCs). Upon ultrasound (US) exposure, cavitation of the ncNBs induces sonoporation, facilitating the delivery of 2’-3’-cyclic GMP-AMP (cGAMP) into the cytosol of APCs. Cytosolic cGAMP activates the stimulator of interferon genes (STING) pathway and downstream signaling, ultimately promoting antitumor activity. *Mouse illustration adapted from NIAID NIH BioArt Source (BIOART-000279)*.

SONATA NBs were engineered for efficient cGAMP loading, CD11b^+^ APC-targeted delivery, and ultrasound-mediated payload release. Here, we demonstrate successful adaptation of the MUSIC-derived functionalization strategy to nanoscale NBs. SONATA enabled ultrasound-guided STING activation in macrophages, enhanced antitumor immunity, suppressed tumor growth in an orthotopic breast-cancer model, and improved therapeutic responses when combined with immune checkpoint blockade. Together, these findings establish SONATA as a systemically administered, ultrasound-responsive immunotherapy platform for spatiotemporally controlled STING activation in cancer.

## Results and Discussion

### Preparation and Characterization of SONATA Platform

To generate an ultrasound-responsive nanobubble platform capable of systemic delivery, we first formulated NBs consisting of maleimide-functionalized phospholipid shells and perfluorobutane (PFB) gas cores. Phospholipids were emulsified with PFB as previously reported [14,15], with modifications to the post-formulation separation process to enrich for nanoscale bubbles while removing larger microbubbles and non-gas-filled particles, including liposomes (Figure 2a). Briefly, the suspension was centrifuged at 50 *g* for 5 min, after which the upper microbubble layer was removed and the remaining infranatant containing NBs was collected via centrifugation at 700 g for 5 min (Figure 1a) for further characterization by nanoparticle tracking analysis (NTA; Particle Metrix, ZetaView).

**Figure 2.**
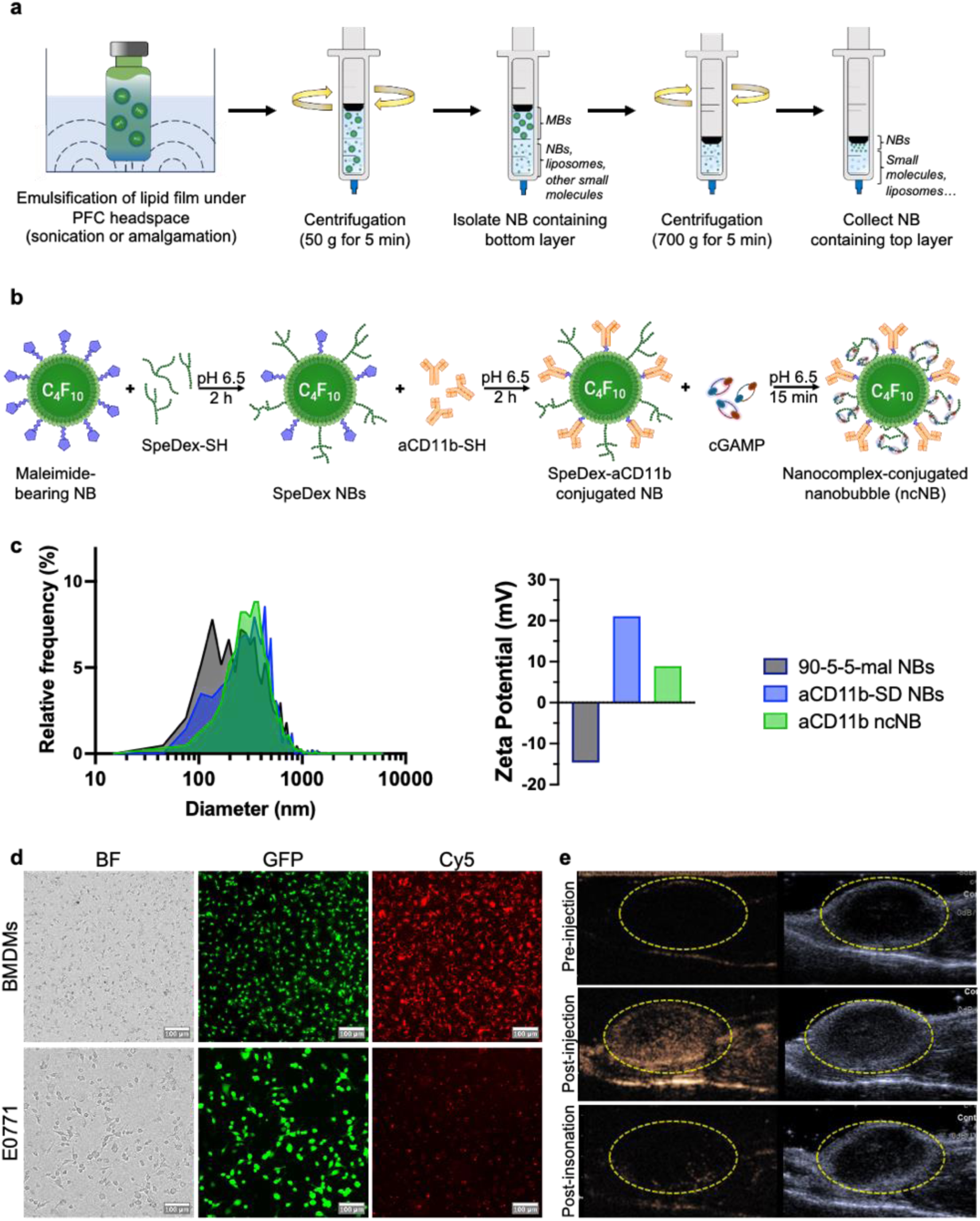
Formulation and characterization of SONATA ncNBs. **(a)** NBs were formulated via emulsification under PFC headspace and then isolated via differential centrifugation. **(b)** Maleimide-bearing NBs were conjugated to SpeDex-SH and aCD11b-SH followed by cGAMP loading. **(c)** Representative ZetaView sizing distribution (left) and DLS zeta potential (right) of NBs after each formulation step. (d) Brightfield and fluorescence microscopy of BMDMs and E0771 cells after incubation with AF647-CD11b-targeted ncNBs validates targeting-specificity. Scale bar = 100 µm. **(e)** ncNBs show enhancement in tumors post i.v injection and collapse upon insonation.

To enable macrophage-directed delivery and enhance cGAMP loading, thiolated anti-CD11b antibody (aCD11b) and cationic spermine-modified dextran (SpeDex-SH) were conjugated to the NB surface through thiol-ene coupling in a one-pot reaction. Following purification, aCD11b-SpeDex NBs were loaded with cGAMP at an N:P ratio of 1:34 to generate cGAMP nanocomplex-loaded NBs (ncNBs) (Figure 2b). All formulations were characterized by NTA and dynamic light scattering (DLS) to determine size distribution, concentration, and surface charge

The resulting ncNBs exhibited a representative mean diameter of 380 nm, a D90 of 590 nm, and a concentration of 5.2 × 10¹⁰ NBs/mL (Figure 2c). Surface charge analysis demonstrated an increase in zeta potential following functionalization with the cationic SpeDex polymer, followed by a decrease after loading the negatively charged cGAMP, consistent with successful nanocomplex formation (Figure 2c).

To evaluate APC-targeted delivery, fluorescently labeled ncNBs were incubated with CD11b⁺ mouse bone marrow-derived macrophages (BMDMs) or CD11b⁻ E0771 breast cancer cells. Brightfield and fluorescence microscopy demonstrated selective targeting of ncNBs to CD11b^+^ BMDMs with minimal binding to CD11b^-^ E0771 cells, consistent with CD11b-mediated targeting of antigen-presenting cells (Figure 2d). Finally, ncNBs retained ultrasound contrast properties and demonstrated tumor enhancement and subsequent collapse following intravenous administration, as visualized using a clinical ultrasound imaging system (Siemens Sequoia with 18H6 linear array transducer), confirming their potential for systemic ultrasound-guided delivery (Figure 2e).

### SONATA enables cGAMP delivery and activation of STING Pathway

To evaluate ultrasound-triggered STING activation by SONATA in vitro, ncNBs were incubated with BMDMs at varying NB-to-cell ratios (200, 250, 500, 750, or 1000 NBs/cell) while maintaining an equivalent cGAMP dose across all treatment groups. Following incubation, ultrasound-mediated activation was performed at 1 MHz, 2 W/cm², with a 50% duty cycle for 60 s. PBS, free cGAMP, and cGAMP-loaded NBs without ultrasound activation were included as control groups.

Ultrasound activation of ncNBs resulted in robust activation of the canonical STING signaling pathway, as demonstrated by increased phosphorylation of STING, TBK1, and IRF3 by Western blot analysis (Figure 3a). Phosphorylation of STING and its downstream effectors TBK1 and IRF3 confirmed activation of the STING signaling cascade, which culminates in IRF3 activation and transcriptional induction of type I interferons. Consistent with activation of this pathway, ELISA analysis of cell culture supernatants revealed significantly elevated IFN-β production in BMDMs treated with ncNBs and ultrasound compared with all control groups (Figure 3b).

**Figure 3.**
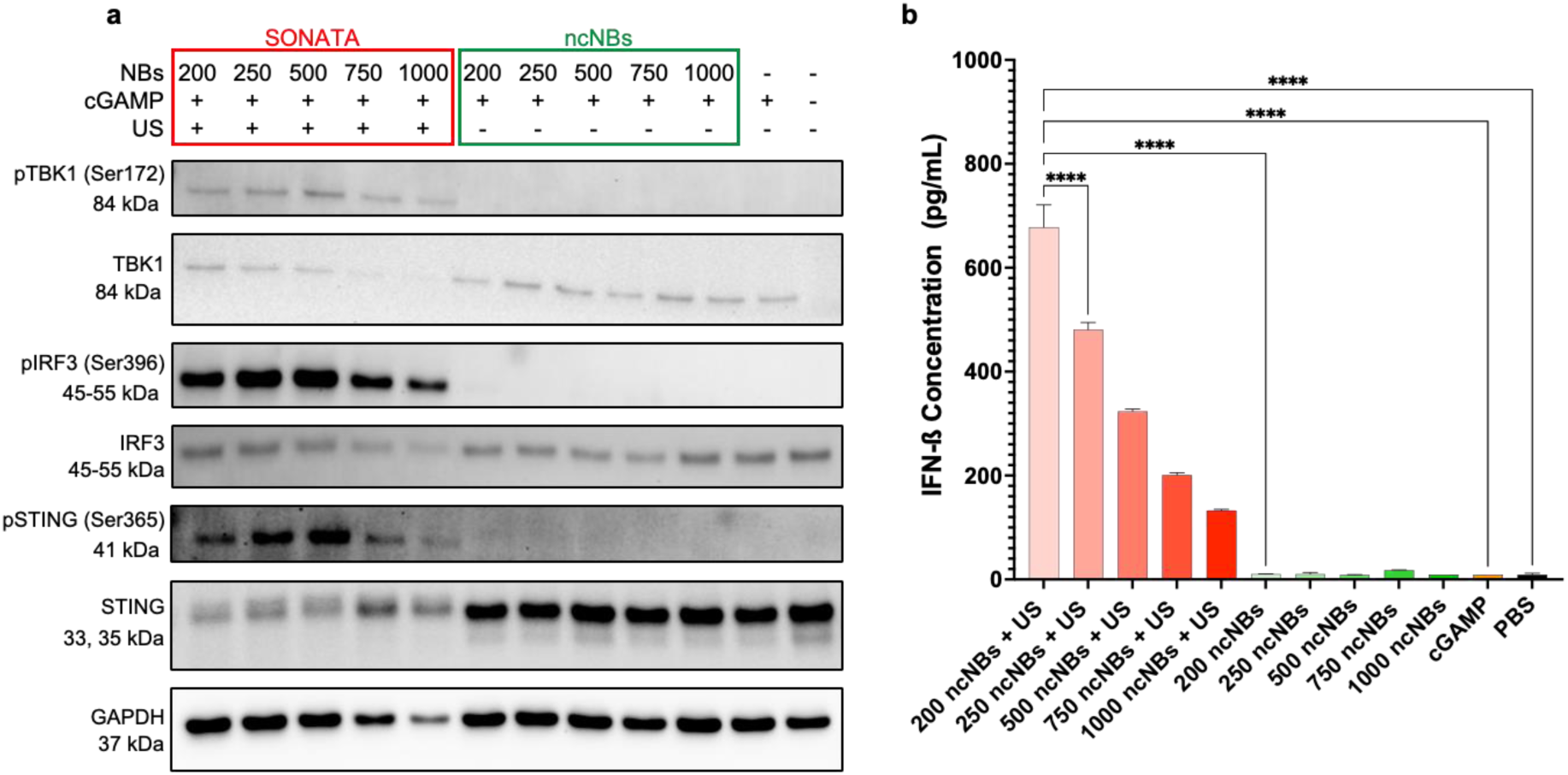
(a) STING activation in vitro with SONATA. BMDMs were treated as indicated followed by **(a)** western blotting to assess phosphorylation of proteins in the STING pathway and **(b)** ELISA for IFN-ß cytokine production 3 h post-treatment. Data were analyzed by one-way ANOVA with Tukey’s multiple comparisons test. p values > 0.05 were considered not significant (ns), p values < 0.05 were considered significant. *p value < 0.05, **p value < 0.01, ***p value < 0.001, ****p value < 0.0001.

Interestingly, despite equivalent cGAMP dosing, IFN-β production decreased with increasing ncNB concentration, with 200 NBs/cell producing the highest response, an approximately 100-fold increase over untreated controls (Figure 3b). The inverse relationship between ncNB concentration and IFN-β production may reflect two formulation-dependent effects. Increasing ncNB concentration at a fixed cGAMP dose increases the N:P ratio, potentially strengthening electrostatic complexation and limiting cGAMP release. In addition, distributing a fixed cGAMP dose across a greater number of ncNBs reduces the cGAMP payload per NB, which may decrease the amount of cGAMP delivered during individual NB–cell interactions. These results demonstrate that SONATA enables efficient ultrasound-dependent cGAMP delivery and activation of the STING–TBK1–IRF3 signaling axis, leading to robust type I interferons production in antigen-presenting cells.

### SONATA activates STING-mediated immune remodeling and inhibits tumor growth

Following demonstration of ultrasound-dependent STING pathway activation in vitro, we evaluated the therapeutic efficacy of SONATA in an immunocompetent orthotopic E0771 breast cancer model. Tumors were allowed to reach 50–80 mm³ prior to treatment initiation, with no significant differences in baseline tumor volumes between groups as determined by ANOVA. Mice received systemic administration of SONATA containing 5 mg/kg cGAMP and 1.39 × 10⁹ NBs/kg via retro-orbital injection. Following systemic administration, tumors were insonated under ultrasound guidance (4 W/cm², 50% duty cycle, 30 s × 3 pulses), with successful injection and acoustic activation monitored using a clinical ultrasound scanner (Figure 2e). Contrast-enhanced ultrasound imaging was used to assess tumor perfusion kinetics and determine optimal timing for ultrasound activation. Treatments were administered every other day for a total of five treatment sessions (Figure 4e).

**Figure 4.**
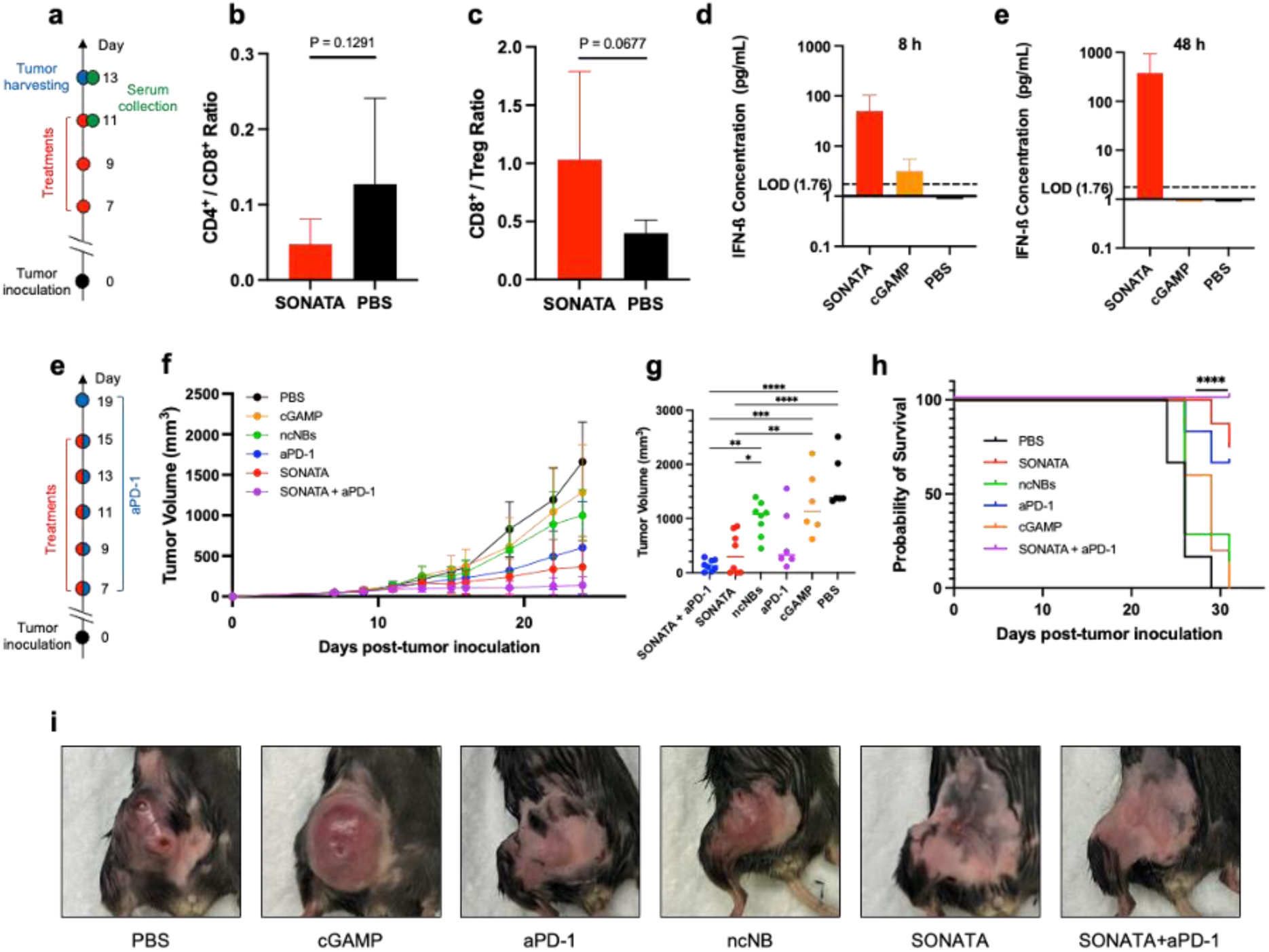
SONATA treatment of E0771 breast cancer. **(a)** Experimental design of SONATA mediated tumor infiltration and IFN-ß production. **b-c** Flow cytometry quantification of tumors 48 hours after the third treatment in SONATA vs. PBS. **(b)** ratio of the number of CD4^+^ T cells to CD8^+^ T cells. **(e)** ratio of the number of CD8^+^ cells to Tregs. **d-e** Accumulation of IFN-ß 8 h **(d)** and **(e)** 48 h post third treatment in tumor determined using ELISA. *n =* 5 replicates for SONATA and *n =* 4 replicates for cGAMP and PBS. **(e)** Treatment schematic for the orthotopic E0771 tumor model. **f–h** Tumor growth and survival were monitored following treatment. **(f)** Tumor growth curves. **(g)** One-way ANOVA comparing tumor volumes on day 24 post-tumor inoculation. **(h)** Kaplan–Meier survival curves for the indicated treatment groups over 31 days. **(i)** Representative images of tumors on day 26 post inoculation. *n* = 6 (aPD-1, cGAMP, PBS) and 8 (ncNBs, SONATA, SONATA+aPD-1). The data represent mean ± s.d. Data were analyzed by one-tailed Welch’s t-test (b,c), one-way ANOVA with Tukey’s multiple comparisons test (f,g) or two-sided log-rank (Mantel–Cox) test (h). p values > 0.05 were considered not significant (ns), p values < 0.05 were considered significant. *p value < 0.05, **p value < 0.01, ***p value < 0.001, ****p value < 0.0001.

To investigate whether SONATA modulates the tumor immune microenvironment (TME), tumors were collected 48 h after the third treatment (Figure 4a) and analyzed by flow cytometry. SONATA treatment resulted in an increased frequency of intratumoral CD8⁺ T cells and a decreased frequency of CD4⁺ T (Figure S3a,b) cells leading to a reduced CD4⁺/CD8⁺ T-cell ratio compared with saline controls (Figure 4b). Additionally, SONATA reduced the frequency of regulatory T cells (Tregs) (Figure S3c), resulting in an increased CD8⁺/Treg ratio (Figure 4c). Although these immune cell changes did not reach statistical significance, they demonstrated a consistent trend toward a more immunostimulatory tumor microenvironment following SONATA treatment. Consistent with localized STING pathway activation, ELISA analysis of tumor lysates revealed detectable IFN-β production only in SONATA-treated tumors, whereas IFN-ß levels remained below limit of detection in control groups 48 h post-treatment (Figure 4e). At 8 h post-treatment, IFN-ß levels were only minimally detectable following cGAMP administration (Figure 4d).

Flow cytometric analysis further demonstrated elevated PD-L1 expression on CD45⁻ tumor cells, providing a rationale for evaluating SONATA in combination with immune checkpoint blockade (Figure S3d). To determine whether SONATA could enhance responsiveness to anti-PD-1 therapy, mice were randomized into six treatment groups following tumor establishment (∼50 mm³): PBS, free cGAMP, non-activated ncNBs, SONATA, anti-PD-1, and SONATA combined with anti-PD-1. SONATA, free cGAMP, and ncNB formulations were administered systemically via retro-orbital injection, while anti-PD-1 was administered intraperitoneally at 10 mg/kg. Ultrasound activation and treatment schedules were performed as described above, with an additional maintenance dose of anti-PD-1 administered 96 h after the final treatment.

Ultrasound-guided activation of SONATA produced significant therapeutic benefit compared with control treatments. In contrast, free cGAMP and non-activated ncNBs did not provide a meaningful survival advantage, demonstrating that both nanobubble-mediated delivery and ultrasound-triggered activation are required for therapeutic efficacy. Combination treatment with SONATA and anti-PD-1 resulted in enhanced tumor growth suppression compared with either treatment alone (Figure 4f-i). This pronounced tumor control was particularly evident during the treatment period and early follow-up (Figure 4f,g).

Long-term survival analysis further demonstrated a therapeutic benefit of SONATA, which increased median survival from 21 to 31 days following randomization compared with PBS-treated mice, corresponding to a 47.6% improvement in median survival (Figure S4g). However, extended monitoring for up to two months revealed eventual tumor progression, indicating that the therapeutic response, although substantial, was not durable under the dosing regimen evaluated. Although SONATA combined with anti-PD-1 showed a trend toward improved survival compared with SONATA monotherapy, this difference did not reach statistical significance (Figure S4g).

Notably, the systemic cGAMP dose used with SONATA was comparable to that previously administered intratumorally with MUSIC. Because only a fraction of an intravenously administered dose is expected to accumulate within the tumor, the effective intratumoral cGAMP exposure achieved with SONATA is likely lower than that achieved by direct intratumoral administration. Thus, optimization of SONATA dose, treatment frequency, and combination scheduling may be required to achieve more durable tumor control.

## Conclusion

In this study, we developed SONATA, a nanobubble-based delivery platform that enables systemic administration and ultrasound-guided delivery of cGAMP to the tumor microenvironment. The nanoscale properties of nanobubbles facilitate systemic circulation and tumor accessibility, while ultrasound provides spatial and temporal control over therapeutic activation. SONATA enhanced antitumor efficacy and demonstrated synergistic therapeutic effects when combined with immune checkpoint inhibition.

Although substantial tumor control was achieved, long-term tumor progression highlights the need for further optimization of systemic SONATA dosing and treatment frequency. In addition, optimization of the timing and dosing of immune checkpoint blockade may further enhance the durability of combination therapy.

This work establishes a versatile biomaterials-based platform that integrates nanoscale delivery with externally controlled activation, offering a promising approach to improve therapeutic precision, potentially reduce off-target effects, and advance the development of next-generation cancer immunotherapies.

## Methods

### NB Formulation

Lipid films containing a mixture of 1,2-distearoyl-*sn*-glycero-3-phosphocholine (DSPC), 1,2-Distearoyl-sn-Glycero-3-Phosphoethanolamine-N-[Methoxy(Polyethylene glycol)-2000] (DSPE-PEG2k), and 1,2-distearoyl-*sn*-glycero-3-phosphoethanolamine-*N*[maleimide(polyethylene glycol)5000] (DSPE-PEG(5000)-mal) with a 90:5:5 molar ratio were prepared. These films were prepared by dissolving DSPC, DSPE-PEG(2000), and DSPE-PEG(5000)-mal in chloroform and slowly evaporating the mixture with a rotary evaporator (Büchi Rotavapor R-100) until mostly dry. The resulting films were then dried overnight under vacuum and stored at −20 °C for later use. The lipid films were solvated in a mixture of PBS 1 ×/propylene glycol/glycerol (80:10:10 v/v/v, 10 mL total) by heating at 70 °C on a heating block for 15 min and sonicated at 70 °C until clear. Perfluorobutane (PFB) vapor was then introduced in the solution, and the resulting mixture was tip sonicated at 70% amplitude for 5 seconds before being cooled down in an ice bath. The resulting NB formulation was centrifuged at low speed (50g, 5 min) to isolate the MB cake and the NB-containing infranatant. NB concentration and size were determined using Nanoparticle Tracker Analysis (NTA, ZetaView equipped with a 520 nm excitation laser, Particle Metrix GmbH, Inning am Ammersee, Germany).

### SpeDex and Anti-CD11b Conjugation onto NBs

Spermine-modified dextran (SpeDex) polymer and thiolated anti-CD11b was prepared as previously reported. Thiolated SpeDex was dissolved in phosphate-buffered saline (PBS) 1 ×, 5 mmol L^-1^ ethylenediaminetetraacetic acid (EDTA, 0.5 ml). at 10 mg mL^-1^ and added to mal-NBs at a 1:20 maleimide:SpeDex molar ratio. The solution was rotated end-over-end for 2 h, followed by addition of thiolated anti-CD11b at 0.05 molar equivalents for an additional 2 h, and then centrifuged at 700g, 5min to isolate the NB cake. The resultant infranatant was re-amalgamated under PFB headspace using Vialmix (Lantheus Medical Imaging) and washed with differential centrifugation (50g, 5 min followed by 700g, 5 min), combined with the previous NB cake and characterized using NTA. To validate targeting, SpeDex-aCD11b MBs were incubated with cGAMP at a nitrogen:phosphate (N:P) ratio of 1:34 and the resulting ncMBs were added to 12-well plates containing either 300,000 mouse bone marrow-derived macrophages (BMDMs) or EO771 murine breast cancer cells. The wells were washed three times, filled with perfluorobutane-saturated PBS, and imaged with bright field and fluorescence microscopy.

### Mice

C57BL/6J mice were bred in-house from founders originally purchased from The Jackson Laboratory and maintained at the animal facility of The University of Texas Southwestern Medical Center or The University of Texas MD Anderson Cancer Center in a specific pathogen free (SPF) environment with an ambient temperature of 22 °C and a relative humidity of 50%. All the mice were maintained on a standard diet and water in a 12 h:12 h light-dark cycle. Tumors were inoculated into the mammary fat pad of 6 to 16 week-old female mice. All animal experiments were approved by and performed in accordance with the Institutional Animal Care and Use Committee of The University of Texas Southwestern Medical Center.

### Cell culture

The mouse mammary carcinoma cell line EO771 was obtained from American Type Cell Culture (ATCC). Mouse bone marrow-derived macrophages (BMDMs) were isolated from the hind leg femur bone marrow of C57BL/6J mice and were cultured or activated with macrophage colony-stimulating factor (M-CSF, 20 ng mL^-1^). EO771 cells and BMDMs were cultured in Dulbecco’s minimum essential medium (DMEM). BMDM media was supplemented with 10% fetal bovine serum (FBS) and 1% penicillin-streptomycin. E0771 media was supplemented with 10% FBS and 1% sodium pyruvate. Cells were incubated at 37 °C under humidified conditions equilibrated with 5% CO_2_. All the cell lines were tested and found to be free of mycoplasma contamination by using MycoAlert kit (Lonza).

### Treatment of BMDMs with cGAMP-Loaded SpeDex-aCD11b NBs

Seven days before treatment, cells were plated at 1,000,000 cells/well in 12-well plates and differentiated using M-CSF. On the day of treatment, SpeDex-aCD11b NBs (200/250/500/750/1000 NBs/cell) were rotated with 10 nmol cGAMP for 15 min to allow binding to yield ncNBs. The medium was aspirated off, and the entire ncMBs volume was added to the wells. Plates were flipped upside down and left to incubate for 10 min at 37 °C. Cells were diluted to 2.5 mL with perfluorobutane-saturated medium, placed on top of a water bath, then sonoporated at 2 W/cm^2^, 50% DC, for 60 seconds using a planar wave transducer (Bulldog-Bio, Sonitron GTS Sonoporation System). cGAMP only, NBs+ultrasound only (NBs), and PBS were used as controls. Cells were collected at 3 or 6 h after treatment for enzyme-linked immunosorbent assay (ELISA) and western blotting.

### Western blotting

Cells were lysed with detergent buffer (50 mM Tris at pH 7.5, 150 mM NaCl, 0.5% Triton-X) with 1X HALT protease inhibitor cocktail (Sigma). Protein samples were collected from the supernatants and loaded on a 4-12% gradient gel (Invitrogen), transferred using iBlot2 to polyvinylidene difluoride (PVDF) membranes (Invitrogen) and blocked in 3% Bovine Serum Albumin (BSA) blocking buffer for 1 h at room temperature. Membranes were then incubated overnight at 4 °C with primary antibodies and then secondary antibodies at room temperature for 1 hour according to the manufacturer’s (Cell Signaling Technology) instructions (Mouse-Reactive STING Pathway Antibody Sampler Kit #16029). The membrane was developed using a ChemDoc MP (BioRad).

### Cytokine assay

Blood samples (∼500 μL) were collected from the orbital sinus of mice that had been anesthetized with isoflurane. Mice were euthanized after blood collection. Samples were then centrifuged at 1500*g* for 15 mins, and the serum was collected. For collection of cell culture supernatants, samples were collected at various times after treatment. Cytokines were measured with the mouse IFN-γ (BioLegend) ELISA Kits according to the manufacturer’s instructions.

### Tumor models

Orthotopic mammary fat pad tumors were established by injecting mouse breast cancer cells (EO771 in C57BL/6J mice at 1 × 10^6^ cells in 100 μL PBS per mouse. Tumors were measured with calipers and tumor volumes were calculated according to an ellipsoid formula (1/2 × length × width^2^). Mice with palpable tumors of similar size were randomized into different groups 7 days after tumor inoculation. Each group consisted of 4-8 mice. Treatments were given on days 7, 9, 11, 13, 15 and consisted of retro-orbital injection (NBs+cGAMP, cGAMP or PBS only in 80-00 μL sterile PBS).

### Ultrasound-guided in vivo STING Activation using ncNBs

Murine breast cancer models were used to test the effectiveness of the nanobubble platform in activating the STING pathway in vivo as follows. Tumors were allowed to grow for 7 days before mice were randomized into three treatment groups (anti-PD-1, SONATA and SONATA + anti-PD-1) and three control groups (PBS only, cGAMP alone, and ncNBs without US activation). The tumors were allowed to grow to 80 mm^3^ and any differences between the means were determined to be non-significant using ANOVA. Tumors were imaged with a clinical US scanner to confirm the successful injection of NBs and peak enhancement in tumors. The mice were injected retro-orbitally with ncNBs suspension given in a bolus injection; the dose was dependent on weight of the mouse, with a dose of 100 μg of cGAMP and 2.7e10 NBs in 100 μL for a 20 g mouse used as the standard. US was applied by using acoustic coupling gel and a 1-MHz plane wave transducer operating at 4 W cm^-2^ for 30 seconds and a 50% duty cycle given to opposite sides of the tumour for a total treatment time of 90 seconds. Mice were treated every two days for a total of five treatments, and tumor size was monitored every other day thereafter. Mice were sacrificed if tumor ulceration appeared or if tumor volume reached 2,000 mm^3^.

### Flow cytometry

The treated mice from each group were euthanized 48 hours post third treatment, and the tumor tissues were collected and digested in 200 U mL^-1^ collagenase IV, 1.6 U mL^-1^ Collagenase I and 15 U mL^-1^ DNase I in 10 mmol L^-1^ RPMI-1640 buffer at 37 °C for 60 min to obtain the cell suspensions. The dissociated cells were then filtered through a 70-μm nylon cell strainer and collected for the following analyses. CD8^+^ T cells, CD4^+^ T cells, and Pan T cells were isolated for flow cytometry analysis as follows. Briefly, cells were first incubated with 1:1000 dilution of LIVE/DEAD Near-IR (Invitrogen cat# L34975) and then blocked with anti-CD16/CD32 (Biolegend Cat # 101320) for 15 min to avoid nonspecific binding to the Fc receptor. Cells were stained separately with different antibodies (anti-CD45-BV510 dilution of 1:160; anti-CD3-PE dilution of 1:80, anti-CD4-AF700; 1:200; anti-CD8-AF488, dilution of 1:200; anti-PD-L1-PE/Dazzle594, dilution of 1:100) according to the manufacturer’s instructions. Cells were then fixed and permeabilized using Biolegend TrueNuclear Transciption Buffer (Biolegend Cat # 424401), after which they were stained with anti-FOXP3-AF647) dilution of 1:50).

### Statistical analyses

All data are shown as means ± standard deviation (s.d.) from at least triplicates conditions unless otherwise indicated. Each experiment was repeated independently at least three times. Statistical analyses included two-tailed Students’s *t*-tests for two groups or one-way analysis of variance with *post hoc* tests for multiple groups, as appropriate. Absorbance data were fitted using a non-linear four-parameter logistic (4PL) regression model. For statistical analysis and graphical presentation, experimental sample concentrations falling below the LOD were treated as censored data and assigned a proxy value of LOD/2 to minimize statistical bias. Survival was determined for mice in every group by the Kaplan-Meier method and compared by the log-rank (Mantel-Cox) test. The *P* values of less than 0.05 were considered to indicate statistical significance. Statistical analyses were done with GraphPad Prism 11 and Microsoft Excel version 11 software. No animals were excluded from the analyses.

