## Supplemental Data for "Systemic Nanobubbles Enable Ultrasound-Guided STING Immunotherapy in Breast Cancer"

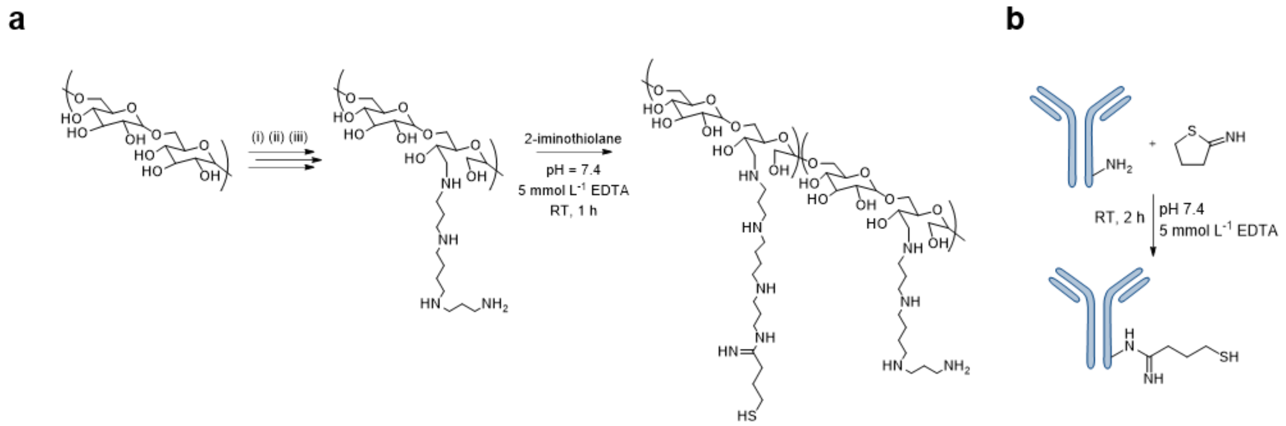


**Figure S1.** Synthesis schemes and characterization of aCD11b. **(a)** SpeDex-SH polymer was synthesized via reductive amination of dextran followed by thiolation with 2-iminothiolane. **(b)** Anti-CD11b antibody was thiolated with 2-iminothiolane to yield aCD11b-SH.

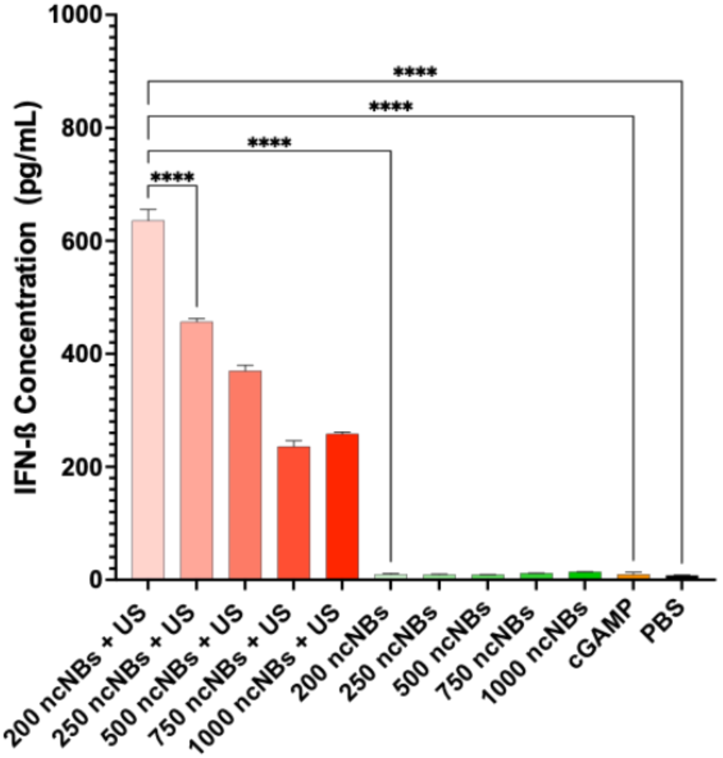


**Figure S2.** ELISA for IFN-ß cytokine production 6 h post-treatment in vitro. Data were analyzed by one-way ANOVA with Tukey's multiple comparisons test. p values > 0.05 were considered not significant (ns), p values < 0.05 were considered significant. *p value < 0.05, **p value < 0.01, ***p value < 0.001, ****p value < 0.0001.

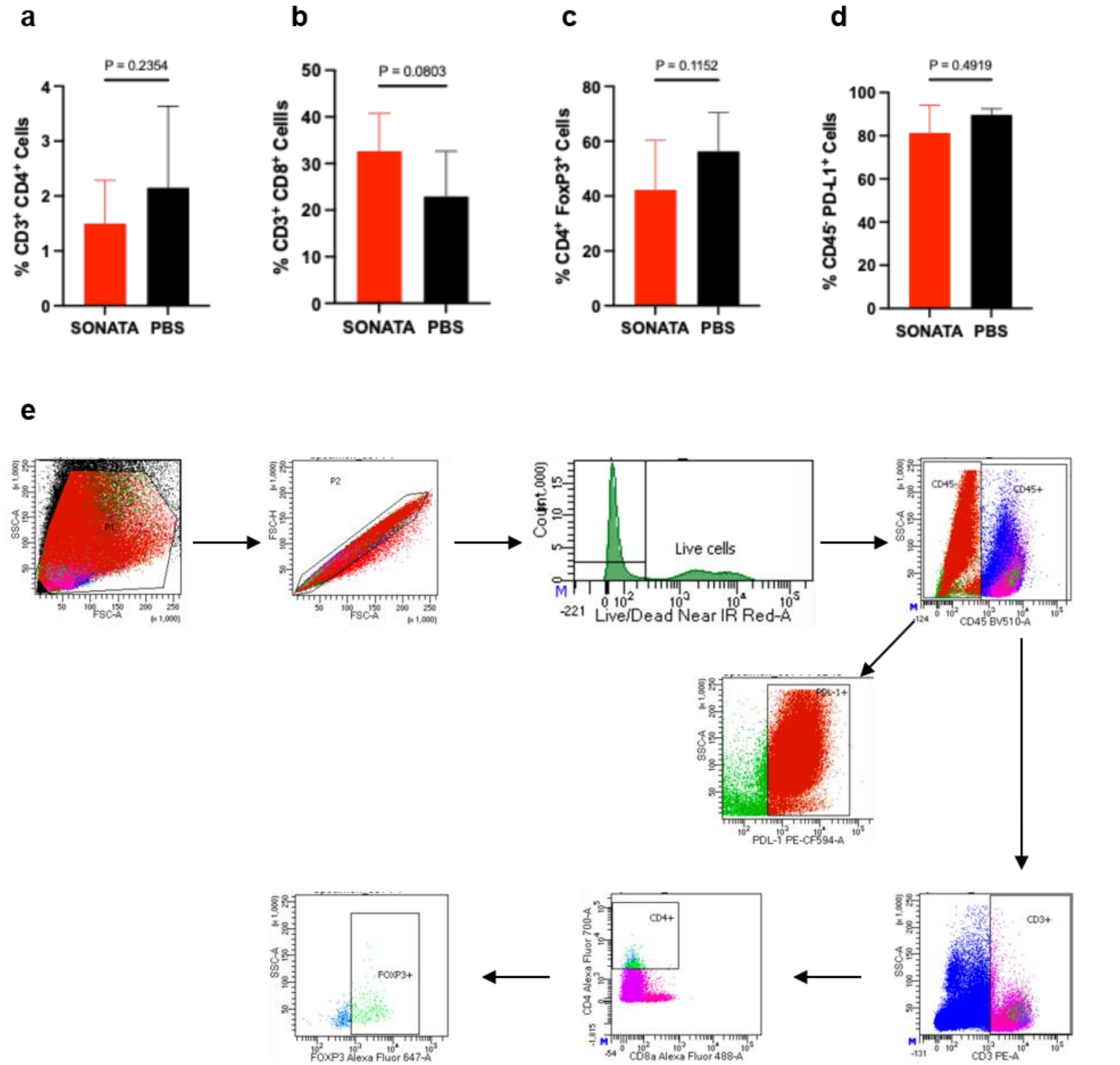


**Figure S3. a-d** Additional flow cytometry quantification of tumors 48 hours after the third treatment in SONATA vs. PBS. Quantification of CD4+ cells **(a)**, CD8+ cells **(b)**, Tregs **(c)**, and PD-L1 in tumor cells **(d)**. **(e)** Flow cytometry gating strategy for **Figs. 4b-c** and **S3**. Data are representative from n=4 (PBS) or n=5 (SONATA) biologically independent experiments. Data are shown as mean ± s.d. and analyzed by Welch's t-test. p values > 0.05 were considered not significant (ns), p values < 0.05 were considered significant.

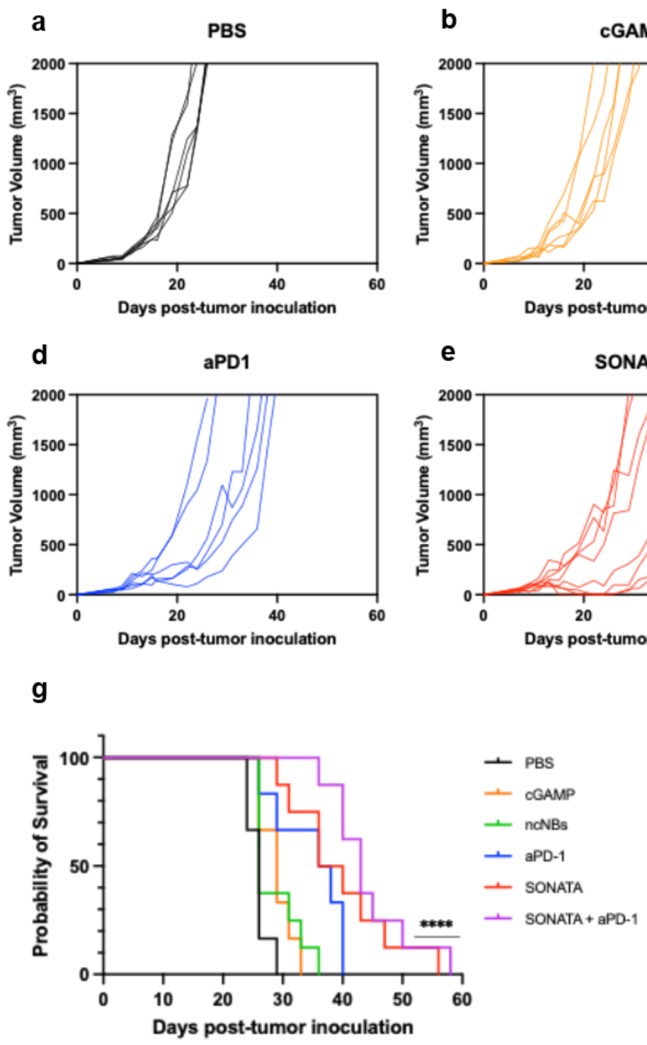


**Figure S4. (a)** Spider plots of individual tumor growth curves. Tumor growth was monitored for 60 days post inoculation or when tumors reached the euthanasia limit (volume over 2000 $mm^3$ or 2 cm in any direction. **(b)** Kaplan–Meier survival curves for the indicated treatment groups over 60 days.
